# Nanoscale organization of Actin Filaments in the Red Blood Cell Membrane Skeleton

**DOI:** 10.1101/2021.03.07.434292

**Authors:** Roberta B. Nowak, Haleh Alimohamadi, Kersi Pestonjamasp, Padmini Rangamani, Velia M. Fowler

## Abstract

Red blood cell (RBC) shape and deformability are supported by a planar network of short actin filament (F-actin) nodes interconnected by long spectrin molecules at the inner surface of the plasma membrane. Spectrin-F-actin network structure underlies quantitative modelling of forces controlling RBC shape, membrane curvature and deformation, yet the nanoscale organization of F-actin nodes in the network *in situ* is not understood. Here, we examined F-actin distribution in RBCs using fluorescent-phalloidin labeling of F-actin imaged by multiple microscopy modalities. Total internal reflection fluorescence (TIRF) and Zeiss Airyscan confocal microscopy demonstrate that F-actin is concentrated in multiple brightly stained F-actin foci ∼200-300 nm apart interspersed with dimmer F-actin staining regions. Live cell imaging reveals dynamic lateral movements, appearance and disappearance of F-actin foci. Single molecule STORM imaging and computational cluster analysis of experimental and synthetic data sets indicate that individual filaments are non-randomly distributed, with the majority as multiple filaments, and the remainder sparsely distributed as single filaments. These data indicate that F-actin nodes are non-uniformly distributed in the spectrin-F-actin network and necessitate reconsideration of current models of forces accounting for RBC shape and membrane deformability, predicated upon uniform distribution of F-actin nodes and associated proteins across the micron-scale RBC membrane.

## Introduction

Plasma membranes of metazoan cells are supported by a thin planar network of short actin filaments (F-actin) cross-linked by spectrin tetramers, forming a membrane skeleton that provides mechanical resilience and creates micron-scale domains of transmembrane proteins such as ion channels and adhesion receptors (Bennett and Baines, 2001; Bennett and Lorenzo, 2013). The spectrin-F-actin network was first discovered in mammalian red blood cells (RBCs), which rely upon the membrane skeleton for their biconcave disc shape and remarkable mechanical deformability and stability, during their transits through the circulation (Chien, 1987; Mohandas and Evans, 1994; Mohandas and Gallagher, 2008). In addition to its mechanical function, the network provides multi-site attachments for RBC transmembrane proteins, including the anion channel (band 3), glycophorin A (GPA) and the glucose transporter (Glut4), among others, restricting their lateral mobility in the membrane (Mohandas and Gallagher, 2008; Kusumi *et al*., 2012b; Alenghat and Golan, 2013; Machnicka *et al*., 2014). Tethering of membrane proteins to the network also facilitates formation of complexes with kinases and/or metabolic enzymes, promoting regulatory signaling and metabolic pathways (Machnicka *et al*., 2014; Lux, 2016). Genetic defects and deficiencies of spectrin-F-actin network components results in abnormal RBC shapes and deformability in human hemolytic anemias (Mohandas and Evans, 1994; Mohandas and Gallagher, 2008; Fowler, 2013; Gallagher, 2013) and affects adaptations of RBCs to invasion by *Plasmodium sp.* in malaria (Dhermy *et al*., 2007; Zuccala and Baum, 2011; Gokhin and Fowler, 2016).

While the molecules and linkages in the RBC membrane skeleton have been mapped in exquisite molecular detail (Burton and Bruce, 2011; Lux, 2016), the nanoscale organization of the spectrin-F-actin network *in situ* across the micron-scale intact RBC is not well understood (Hoffman, 2001; Fowler, 2013; Lux, 2016). The current understanding of the RBC spectrin-F-actin network is based on negative staining electron microscopy (EM) of spread membrane skeletons in which (α1β1)_2_–spectrin tetramers are extended to their full ∼200 nm length (Fowler, 2013; Gokhin and Fowler, 2016; Lux, 2016). These spread images reveal a periodic 2D quasi-hexagonal lattice of short ∼37 nm F-actin nodes connected by ∼200 nm long (α1β1)_2_–spectrin tetramers, with 5-7 spectrins attached to each F-actin node (Byers and Branton, 1985; Shen *et al*., 1986; Liu *et al*., 1987). With a native membrane surface area of 140 μm^2^ and ∼35,000 F-actin nodes (junctional complexes) in the membrane skeleton of each RBC (Fowler, 2013; Lux, 2016), simple calculations reveal that the spread network of extended (α1β1)_2_-spectrin tetramers would encompass a surface area ∼9-fold greater than the cell membrane. Thus, in the native RBC membrane, the 2D lattice is condensed, with quaternary folding of spectrin-repeat domains believed to provide network extensibility for the membrane deformations essential for the survival of the cell in the circulation (McGough and Josephs, 1990; Ursitti *et al*., 1991; Terada *et al*., 1996; Nans *et al*., 2011; Brown *et al*., 2015). In a resting biconcave RBC, the network has been presumed to be uniformly condensed across the entire micron-scale membrane area of the cell, with spectrin tetramers all ∼1/3 their extended length, so that F-actin nodes are on average ∼70 nm apart. A presumption of uniform network dimensions in resting RBCs has formed the basis for many quantitative models of forces to account for the biconcave shape, and membrane deformations under flow or mechanical perturbations (Discher *et al*., 1998; Li *et al*., 2005; Li *et al*., 2007; Fedosov *et al*., 2010; Peng *et al*., 2013; Li and Lykotrafitis, 2014; Chen and Boyle, 2017; Fai *et al*., 2017).

A periodic spacing of F-actin nodes and their connecting spectrin strands has been supported by super-resolution fluorescence microscopy and autocorrelation analysis of F-actin and associated proteins, showing F-actin nodes uniformly spaced ∼80 nm apart (Pan *et al*., 2018). By contrast, previous studies visualizing the spectrin network in the unspread membrane skeleton by electron microscopy (EM) and atomic force microscopy approaches revealed a dense, irregular meshwork of short spectrin strands with a wide range of strand lengths, from ∼20 to ∼80 nm (Ursitti *et al*., 1991; Terada *et al*., 1996; Takeuchi *et al*., 1998; Swihart *et al*., 2001; Nans *et al*., 2011). While F-actin nodes could not be identified due to network density and the presence of associated proteins obscuring filament structure, a cryo-electron microscopy study of Triton-extracted membranes was able to identify short F-actins based on their connections to multiple, highly convoluted spectrin strands of widely varying lengths, 46 +/- 15 nm, again implying a non-uniform network organization (Nans *et al*., 2011). Using the varying spectrin strand length data from this study, as well as new proteomics data showing increased spectrin density in RBCs (Bryk and Wisniewski, 2017; Gautier *et al*., 2018), a quantitative model has been developed recently which provides a better explanation of the observed physical behavior of the membrane during RBC membrane deformations (Feng *et al*., 2020).

A further complexity is that current structural models of the periodic spectrin-F-actin lattice in RBCs are predicated upon the idea that the short F-actin nodes are equivalent and function as stable linkages in unstressed biconcave RBCs. This assumption has been challenged by observations that RBCs contain cytosolic actin monomers (G-actin) at concentrations sufficient to allow barbed end assembly, comprising about 5% of total RBC actin (Gokhin *et al*., 2015). In addition, a subset (∼10%) of the membrane skeleton F-actin can assemble or disassemble, based on treatment of RBCs with jasplakinolide (Jasp) or Latrunculin A (LatA), respectively. RBC membrane mechanical properties are influenced by F-actin dynamics since perturbation of the G/F-actin ratio by actin drugs affects membrane tension and leads to increased cell deformability (Gokhin *et al*., 2015; Gokhin and Fowler, 2016). The existence of an F-actin subpopulation with dynamic properties raises the possibility that F-actin organization may be inhomogeneous across the RBC membrane.

Here, we examined directly the nanoscale distribution of F-actin in intact RBCs using fluorescent-phalloidin labeling of cells imaged by Total Internal Reflection Fluorescence (TIRF) microscopy (Mattheyses *et al*., 2010), super-resolution Zeiss AiryScan confocal microscopy (Huff, 2015) and Stochastic Optical Reconstruction Microscopy (STORM) (Xu *et al*., 2013; Pan *et al*., 2018). Our results demonstrate that F-actin appears in a patchy distribution with brightly staining F-actin foci interspersed with dimmer F-actin staining regions, which are observed to rearrange dynamically. Single molecule STORM imaging and computational cluster analysis of experimental and synthetic data sets indicates that the bright foci correspond to groups of filaments with single filaments located between clusters. This implies that the spectrin-F-actin network is not homogeneously organized, and instead that some F-actin nodes are likely connected by un-extended (α1β1)_2_–spectrin tetramers forming groups of filaments, while other F-actin nodes are connected by extended tetramers, forming solo filaments. These data will necessitate reconsideration of current molecular, structural and mechanical models of RBC shape and membrane deformations predicated upon the idea that the F-actin nodes and their associated membrane proteins are uniformly distributed across the membrane.

## Results

### F-actin is non-uniformly distributed across the membrane

To address the organization of the F-actin network in RBCs *in situ*, we imaged F-actin by fluorescent-phalloidin labeling of fixed and permeabilized human RBCs under conditions that preserve cell integrity (Smith *et al*., 2018). Epifluorescence images focused on the edge of a biconcave cell double-labeled with rhodamine-phalloidin (rho-phalloidin) and an antibody to the membrane marker glycophorin A (GPA) reveal uniformly distributed, bright rim staining corresponding to the membrane at the edge of the biconcave disc (Figure 1A). A smaller fuzzy circle near the center of the cell corresponds to the tangentially oriented membrane at the edge of the dimple. Due to the superimposition of the out-of-focus light in epifluorescence microscopy, it is impossible to obtain more detailed information regarding the lateral distribution of F-actin in the plane of the membrane.

**Figure 1.**
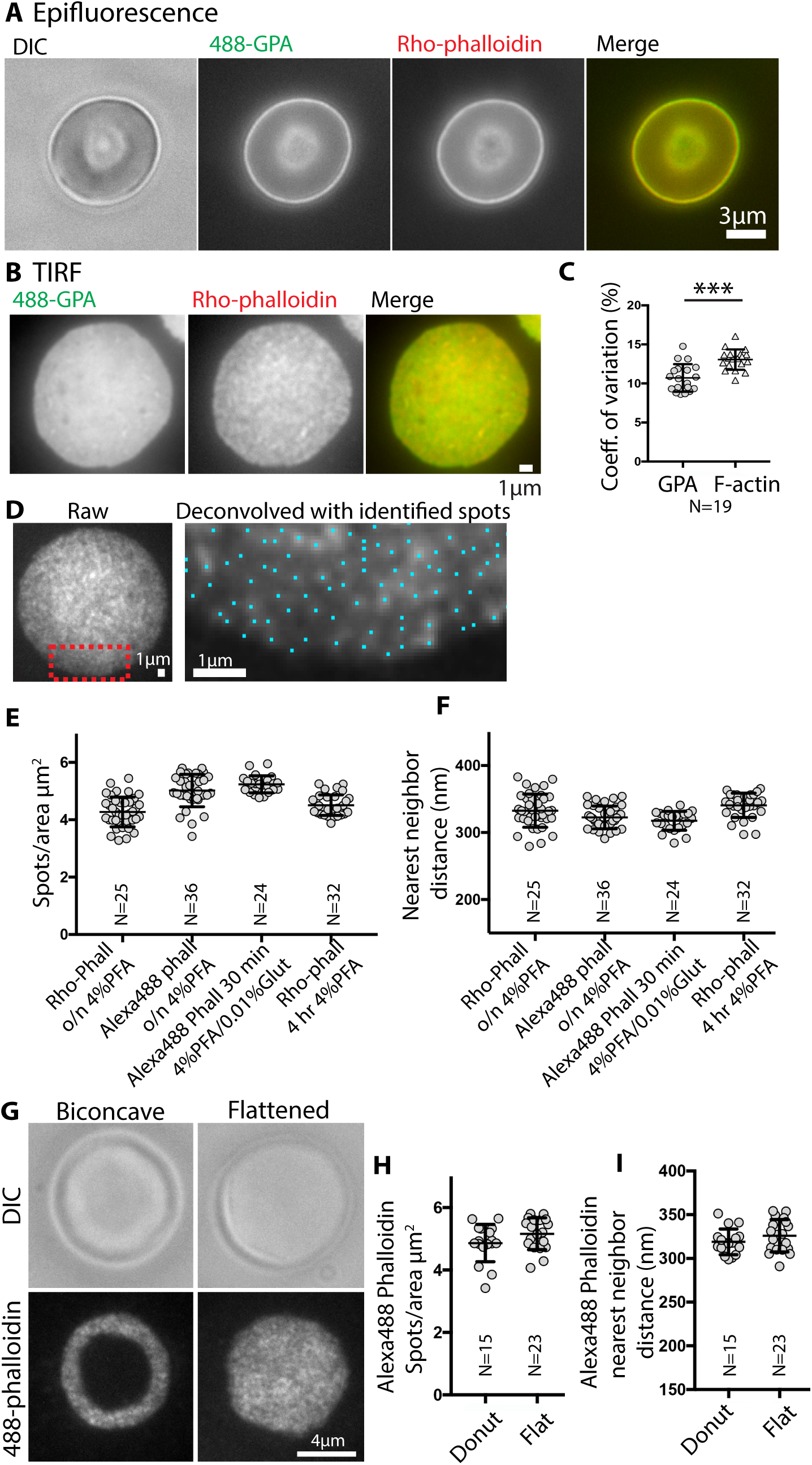
TIRF microscopy reveals bright F-actin foci at the RBC membrane. (A) Differential interference contrast (DIC) and epifluorescence microscopy images of a biconcave RBC stained with Alexa 488-anti GPA antibodies (488-GPA) (green) and Rhodamine (Rho)-phalloidin (red); merge on far right. Bar, 3 μm. (B) TIRF images of a flattened RBC stained with Alexa 488-GPA (green) and Rho-phalloidin (red); merge on far right. Bar, 1 μm. GPA staining is relatively smooth and continuous, compared to uneven F-actin staining. (C) Coefficients of variation for GPA and F-actin staining intensities, determined from images as in B. p <0.0001. N = 19 cells. (D) Left, unprocessed TIRF image of a flattened RBC stained with Rho-phalloidin. Right, higher magnification deconvolved image of portion of cell in red box in left panel. Blue dots, centroids of F-actin foci determined in Nikon Elements. Bars, 1 μm. Quantification of (E) F-actin foci density (spots/μm^3^) and (F) nearest neighbor distance (nm) from high magnification deconvolved images of RBCs stained with Rho- or Alexa-488 phalloidin. No significant differences are observed for RBCs stained with different phalloidins or fixed under different conditions indicated. Each point represents data from an individual cell. (G) DIC or TIRF microscopy images of a biconcave or flattened RBC stained with Alexa 488-phalloidin (488-phalloidin). Bar, 4 μm. (H) Quantification of F-actin foci density (spots/μm^3^) and (F) nearest neighbor distance (nm) from high magnification deconvolved images of biconcave or flattened RBCs stained with Alexa 488-phalloidin. Each point represents data from an individual cell. No significant differences are observed between biconcave and flattened RBCs.

Therefore, we turned to TIRF microscopy, which selectively illuminates fluorophores within 100-200 nm from the coverslip, at or near the cell membrane (Mattheyses *et al*., 2010). Since the membrane surface area visible in TIRF images of biconcave-shaped RBCs is limited, we first examined fixed cells that had flattened during cytospinning onto the coverslip (113 *x g*). Comparison of F-actin with GPA staining in these flattened cells reveals an irregular, patchy appearance of F-actin staining contrasting with the smooth, continuous GPA staining across the membrane (Figure 1B). The irregular pattern for F-actin staining is also evident from the significantly greater coefficient of variation for F-actin pixel intensities as compared to GPA for each cell (Figure 1C). Following deconvolution, at higher magnification, the uneven F-actin staining appears as spots or foci of varying brightness, whose centroids can be identified using the General Analysis tools in Nikon Elements image analysis software (Figure 1D). Quantification reveals a density of ∼4-5 spots/μm^2^, and an average nearest neighbor distance of ∼300-350 nm (Figure 1E, F).

To rule out the possibility that fixation conditions or loss of biconcave RBC shape may induce artifactual rearrangements and clustering of F-actin, we compared several fixation conditions and biconcave *vs.* flattened cells. The irregular F-actin staining pattern, spot density and nearest neighbor distances are similar for RBCs fixed with 4% PFA overnight or for 4 hours, or with 4% PFA plus 0.01% glutaraldehyde for 30 minutes (Figure 1E, F). We also compared rho-phalloidin to Alexa 488-phalloidin labeling and observed a similar irregular F-actin staining pattern for both probes (Figure 1E, F). While the spot density tended to be slightly less, and nearest neighbor distances correspondingly slightly greater for rho-phalloidin than Alexa 488-phalloidin, this did not achieve statistical significance (Figure 1E, F). These small differences may be because Alexa 488-phalloidin staining of RBC F-actin is brighter than that of rho-phalloidin, enabling better detection of more closely spaced, dimmer fluorescence spots. The irregular F-actin staining pattern, spot density and nearest neighbor distances are similar for fixed RBCs that remained biconcave or were flattened mechanically during deposition on the coverslip by cytospinning (Figure 1G-I). These experiments demonstrate that a patchy F-actin distribution (F-actin foci) along the membrane is a robust feature of RBCs.

### F-actin foci are exclusively associated with the membrane

To further explore F-actin distribution on the membrane at higher resolution and to rule out potential contributions from F-actin in the cytoplasm, we used sensitive Zeiss AiryScan confocal fluorescence microscopy to image GPA and F-actin staining in fixed biconcave RBCs. This method has an XY resolution of ∼140 nm and XZ resolution of ∼300 nm (Huff, 2015), somewhat better than the maximum resolution of TIRF microscopy (∼200 nm) (Mattheyses *et al*., 2010). Maximum intensity projections of Z stacks of RBCs stained with Alexa 488-conjugated anti-GPA revealed smooth and continuous GPA staining along the membrane. By contrast, in separate cells, the Alexa 488-phalloidin staining for F-actin was clearly discontinuous and patchy (Figure 2A). We also double-labeled cells with Alexa 488-anti-GPA and rho-phalloidin, or with Alexa 488-phalloidin and rhodamine-Wheat Germ Agglutinin (WGA), which labels GPA, the major sialylated membrane protein (Figure 2B). In RBCs with an orthogonal orientation that had attached to the coverslip at their rims, single XY optical sections through the middle of the cells demonstrated that the GPA (or WGA) and F-actin staining were exclusively associated with the membrane, and with a globally similar intensity at the rim as compared to the dimple (Figure 2B). Due to the oversampling of Z slices and the Z stretch, it is not possible to evaluate the F-actin distributions in the plane of the membrane from these optical sections through the middle of the cell.

**Figure 2.**
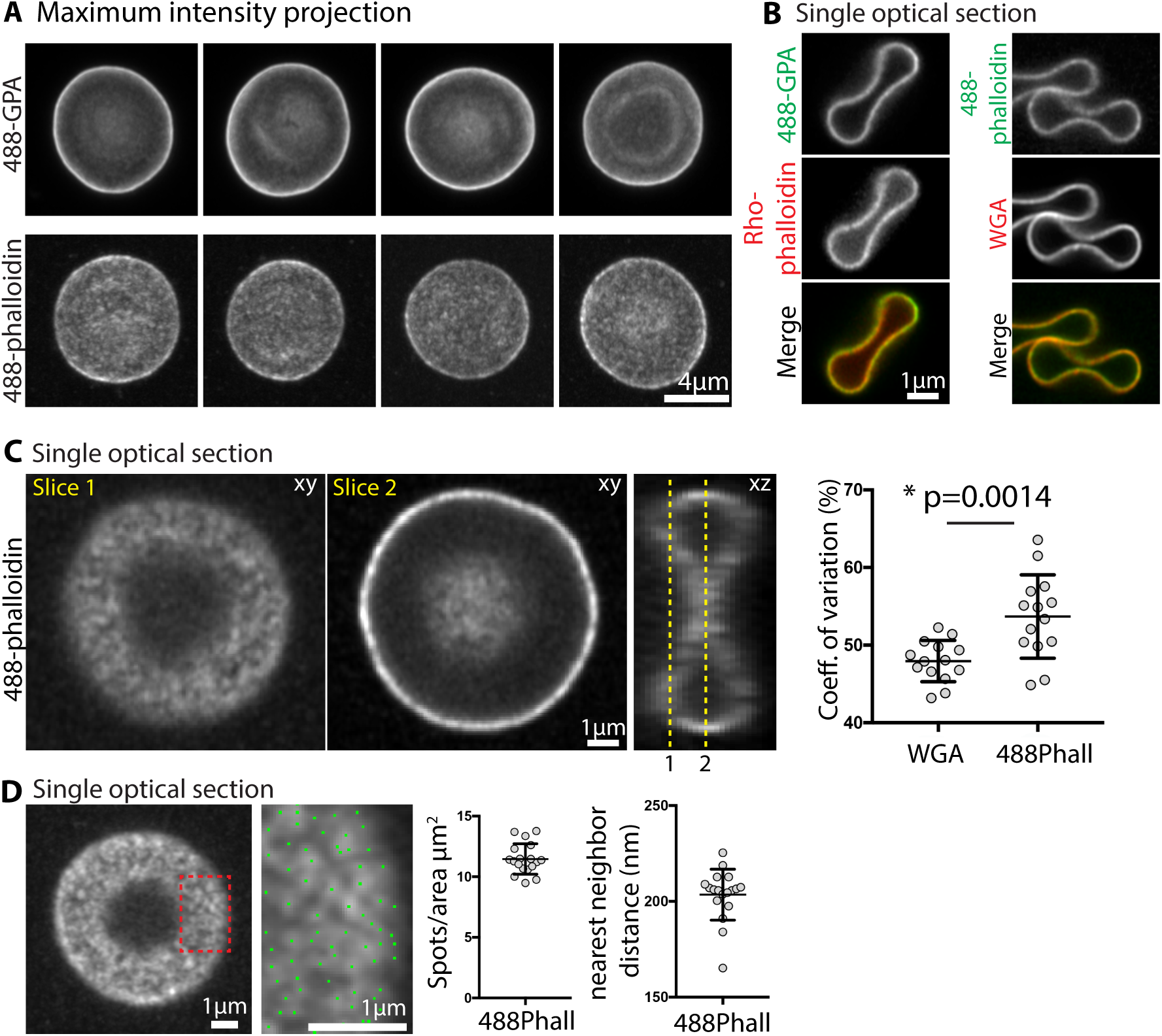
Superresolution Zeiss AiryScan confocal fluorescence microscopy reveals bright F-actin foci at the RBC membrane. (A) Maximum intensity projections of an AiryScan confocal Z-stack of RBCs stained with Alexa 488-antibodies to GPA (488-GPA) or with Alexa-488 phalloidin for F-actin. GPA staining is smooth and continuous, while F-actin staining is irregular and patchy. Bar, 4 μm. (B) Single XY optical sections from Z stacks of RBCs stained with Alexa 488-GPA and Rho-phalloidin (left vertical panels), or with Alexa 488-phalloidin and rhodamine-wheat germ agglutinin (WGA) (right vertical panels). Merges at bottom. Bar, 1 μm. Images of the membrane in cross-section at the rim and dimple regions of cells attached end-on to coverslip show that F-actin staining is exclusively associated with the membrane, colocalizing with GPA or WGA, with similar intensity at the dimple and rim. (C) Single XY or XZ optical sections from Z stacks of RBCs attached parallel to the coverslip, stained with Alexa 488-phalloidin. Left, XY optical section immediately adjacent to the coverslip, revealing irregular and patchy F-actin staining along the membrane (Slice 1). Middle, XY optical section through the middle of the cell revealing bright uneven F-actin staining along the membrane at the rim. Bar, 1 μm. Right, XZ optical section from the middle of the same cell, oriented orthogonally to the XY section in Slice 2. Membrane curvature and oversampling of fluorescence in optical sections leads to apparently continuous F-actin staining along the rim in the XY image in Slice 2. This precludes accurate evaluation of F-actin distribution in cross-sectional views of the RBC membrane in either XY or XZ views. Right graph, coefficient of variation for Alexa-488 phalloidin staining at the membrane is significantly higher than for rhodamine-WGA staining of GPA. Each point represents data from an XY optical section at Slice 1 of an individual cell. p < 0.0014. N = 14. (D) Left image, single XY optical section from a confocal Z-stack of RBC stained with Alexa 488-phalloidin. Right image, higher magnification image of portion of cell in red box in left panel. Blue dots, centroids of F-actin foci determined in Nikon Elements. Bars, 1 μm. Right graphs, Quantification of F-actin foci density (spots/μm^2^) and nearest neighbor distance (nm) from high magnification single optical sections from images as shown on left. Each point represents data from an individual cell. N = 16 cells.

Therefore, we examined single XY optical sections from the rim region of the RBC at the coverslip, in the plane of the membrane, which again revealed an irregular pattern of Alexa-488-phalloidin F-actin staining intensity across the RBC membrane (Figure 2C, Slice 1). Similar to TIRF images, the coefficient of variation of pixel intensities was significantly greater for the F-actin than for the WGA (Figure 2C). A patchy distribution of F-actin is also detected in the dimple region in an optical section through the middle of the cell (Figure 2C, Slice 2), but due to the curvature at the edges of the dimple it is difficult to identify discrete foci for analysis. Quantitative analysis of F-actin distributions by determination of the centroid positions for F-actin foci at the rim region showed about ∼12 spots/μm^2^ with a nearest neighbor distance of ∼200nm (Figure 2D). In these AiryScan images, the F-actin spot density is about two to three times greater, and the nearest neighbor distances are about half of those determined from the TIRF images at the rim region (Figure 1G-I). This can be ascribed to the increased XY resolution of the AiryScan *vs*. TIRF (∼140nm *vs*. ∼200 nm), as well as to a greater sensitivity of the AiryScan fluorescence image detection compared to TIRF imaging, likely allowing discrimination of more closely spaced, dimmer spots. In summary, the irregular F-actin staining distribution across the RBC membrane observed with both imaging modalities indicates that a large proportion of the RBC F-actin in the spectrin-F-actin network is present in small regions, or foci, exclusively associated with the RBC membrane.

### F-actin foci dynamically rearrange in live cells

Next, we wondered whether F-actin foci are present in live RBCs and whether they are stable features of the spectrin-F-actin network over time. To address this, we imaged live RBCs labeled with a cell-permeable, infrared-fluorescent Jasplakinolide (Jasp) conjugate, referred to as SiR-actin (SiR-actin) (Lukinavicius *et al*., 2013; Gokhin *et al*., 2015). We used TIRF microscopy to visualize F-actin foci at the RBC rim closest to the coverslip (Figure 3A, B), and a Leica-SP8 Hyvolution system to visualize F-actin foci in the dimple region of biconcave RBCs (Figure 3C). Both the rim and dimple regions of live RBCs show a patchy distribution of SiR-actin fluorescence intensity, similar to that observed at the rim region at the membrane of fixed RBCs imaged by TIRF microscopy. Visual inspection of successive time-lapse images collected for intervals of 2 seconds revealed appearance and disappearance of some foci with others persisting for several frames (Figure 3B, C). Some of the individual F-actin foci appear to undergo lateral movements back and forth, but do not undergo long-range directed translocation or flow. We could not further extend these qualitative observations, since we were unable to image the SiR-actin in live RBCs for shorter time intervals, or for more than a total of 10 sec, due to extensive photobleaching. Nevertheless, these observations in living cells indicate that the F-actin foci are dynamic structural features of the F-actin network *in situ*.

**Figure 3.**
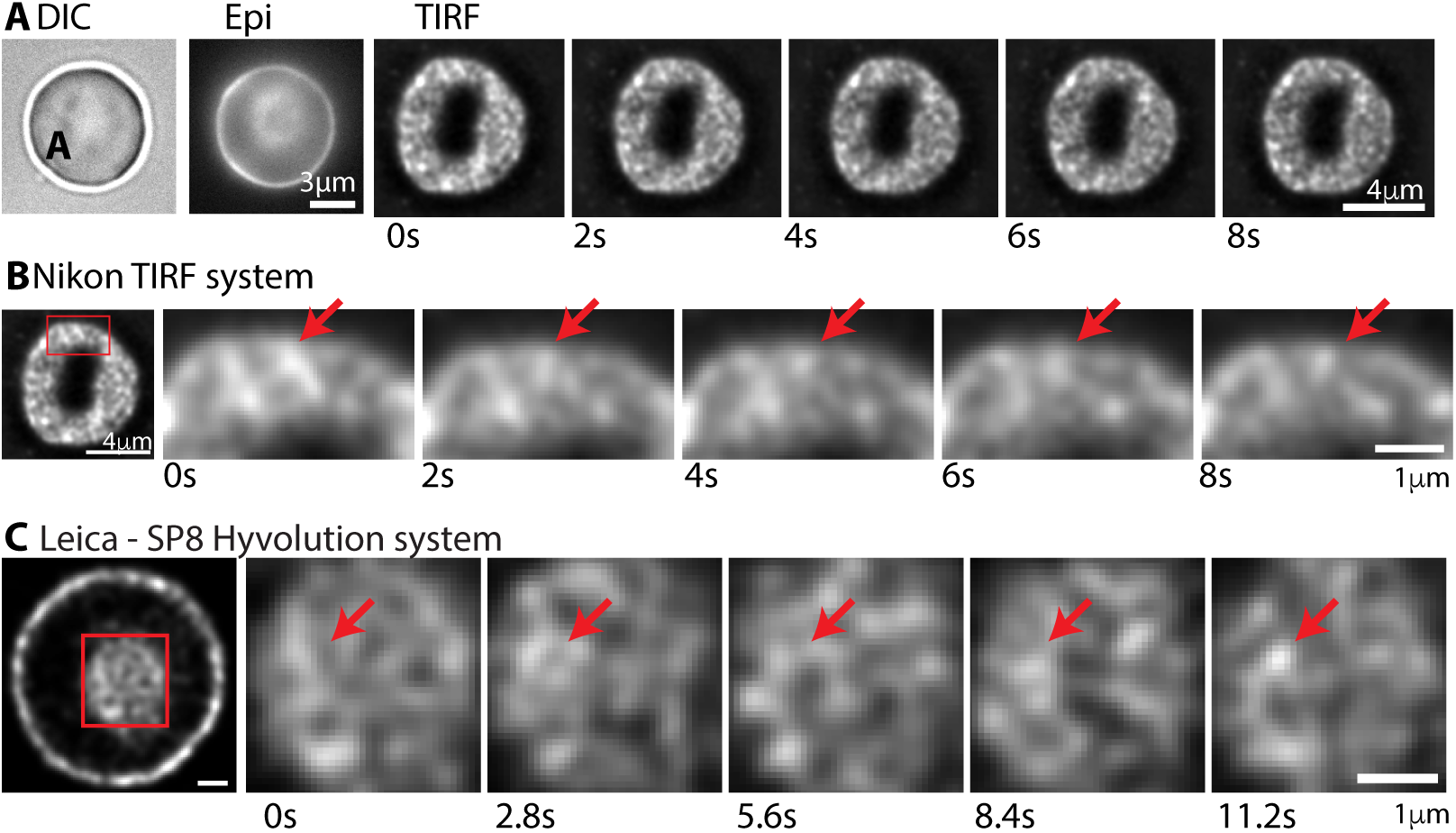
Time-lapse imaging of F-actin foci in live RBCs reveals dynamic movements. (A) DIC and epifluorescence images of a SiR-actin-labelled biconcave RBC before collection of sequential TIRF microscopy images of the same RBC for 2 sec intervals, over 10 sec. Bar, 4 μm. (B) Higher magnification images of a portion of the same RBC. Changes in appearance or positions of F-actin foci are indicated by red arrows. Bar, 1 μm. (C) Images of a SiR-actin-labelled biconcave RBC acquired by Leica-SP8 Hyvolution microscopy (see Methods). Far left panel, XY optical section near the middle of the cell, revealing F-actin foci along the membrane at the rim, and at the membrane of the dimple region (box). Right panels, sequential images of the dimple region collected for 2.8 sec intervals, over 10 sec. The dim fluorescence of the SiR-actin in the RBCs as well as photobleaching precluded collecting more images for shorter time intervals, or over a longer time period. Bar, 1 μm.

#### Single molecule TIRF/STORM imaging demonstrates non-uniform distribution of F-actins

To further analyze the nanoscale F-actin organization in the spectrin-F-actin network, we performed TIRF microscopy of RBCs stained with Alexa 647-phalloidin and imaged under conditions suitable for STORM (Xu *et al*., 2013; Pan *et al*., 2018). We acquired ∼100,000 frames from each sample and used Nikon Elements software to obtain single-molecule localizations of Alexa 647-phalloidin-labeled F-actins, revealing a map of molecular localizations across the entire RBC surface (Figure 4A), with a localization precision of ∼20 nm (Supplemental Figure S1). The Alexa 647-phalloidin molecular localizations reconstructed from STORM images appeared distributed unevenly across the cell surface (Figure 4A), consistent with TIRF images (Figure 1), and AiryScan images (Figure 2).

**Figure 4.**
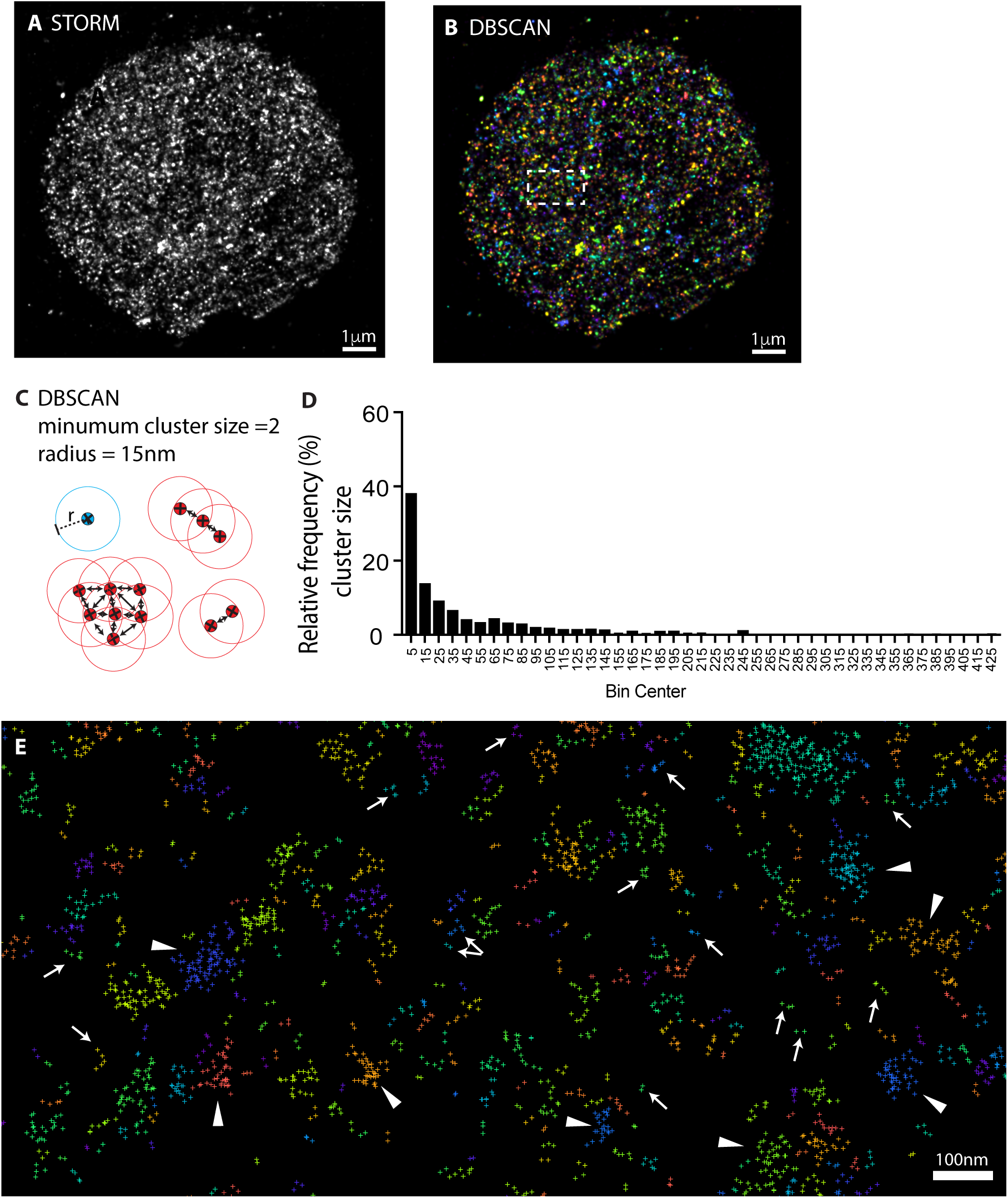
TIRF/STORM imaging and DBSCAN analysis reveals a clustered distribution of individual Alexa 647-phalloidin molecules indicative of single F-actins interspersed with groups of multiple F-actins at the RBC membrane. (A) STORM image of Alexa 647-phalloidin molecules in an RBC, acquired as in Methods. (B) DBSCAN analysis of cluster formation by Alexa 647-phalloidin molecules, with individual Alexa 647 molecules in the same cluster color-coded the same color, and adjacent clusters different colors. White dotted box shown at higher magnification in panel E. Bar, 1 μm. (C) DBSCAN parameters used to define classification of Alexa 647-phalloidin molecules in a cluster. The minimum cluster size was set to 2, and molecules are included in a cluster only if they are within 15 nm. (D) Histogram of cluster sizes and frequency, showing a range of cluster sizes from 2-10 molecules all the way up to >400 molecules, with smaller clusters more abundant than larger clusters. Cluster sizes are grouped into bins of 2-10, 11-20, etc., and centered on 5, 15, etc., for purposes of display. (E) High magnification image of boxed region in panel B, showing a visual depiction of clusters determined by DBSCAN. Molecules in the same cluster are the same color, while adjacent clusters are a different color. Arrows, clusters with 2-10 Alexa 647-phalloidin molecules, often arranged linearly and likely representing individual filaments. Arrowheads, clusters with 11 or greater Alexa 647-phalloidin molecules, likely representing a group of filaments. Bar, 100 nm.

We utilized the density-based spatial clustering of applications with noise (DBSCAN) algorithm embedded in Nikon Elements software to analyze molecular clustering of the Alexa 647-phalloidin signals across the entire cell surface (Figure 4B). In this analysis, molecules are considered part of a cluster if they have a number of neighbors equal to or greater than a minimum number, for a specified radius, r (Figure 4C) (Pedregosa *et al*., 2011; Schubert *et al*., 2017). We selected an r value based on the minimum detectable distance between two points. With a spot size of 20.68 nm +/- 3.77 nm to 22.63 nm +/- 5.83 nm (S.D.) for individual cells (Supplemental Figure 1A) and using FWHM = 2.355 σ (where σ is the standard deviation), we calculated the FWHM to be between 8.88 nm and 13.73 nm (Supplemental Figure 1B). Thus, we selected a radius of 15 nm for the minimum distance at which two molecules could be distinguished and form a cluster.

Using a minimum cluster size of 2 and a radius of 15 nm we identified a wide range of cluster sizes, containing from 2 up to 650 molecules, with most of the Alexa 647-phalloidin molecules in small clusters of <20 molecules, and the remainder in larger clusters ranging from 20 up to >200 molecules (Figure 4D). To visualize the locations of clusters, each cluster of molecules was assigned a different color (Figure 4B, E). Small clusters of 2-10 molecules are visible throughout the field (Figure 4E, arrows), interspersed with larger clusters of 20 or more (Figure 4E, arrowheads). The smallest clusters with 2-10 Alexa 647 molecules are often linearly arranged, with a length <50 nm, suggesting they could represent individual short F-actins, 37nm +/- 5nm long (refs). Following this reasoning, the larger clusters with up to ∼300 Alexa 647 molecules could contain up to ∼30 short filaments (assuming ∼10 Alexa 647-phalloidin-labeled actin subunits/filament). These large clusters are irregularly shaped and can be >100 nm across, and likely correspond to groups of multiple filaments in close proximity to one another (Figure 4E). To further explore the distributions of small and large clusters, we visualized only the small clusters (2-10) (Figure 5B, D, D’) or the larger clusters (11 and above) (Figure 5C, E, E’). Pseudo-coloring small and large clusters sizes in green and red, respectively, shows clearly that single F-actins (defined as clusters of 2-10 Alexa 647-phalloidin molecules) are located between larger clusters containing multiple filaments (defined as clusters ≥11 molecules) (Figure 5F, F’). Quantitative analysis shows that approximately 40% of Alexa 647 molecules are associated with small clusters (2-10) likely representing single filaments, while 60% are in larger clusters (≥11), containing multiple filaments (Figure 5G). We conclude that somewhat more than half of all F-actin is present in large groups of multiple filaments, with slightly less than half of the F-actin in the RBC membrane skeleton as solitary filaments, located in between the larger groups.

**Figure 5.**
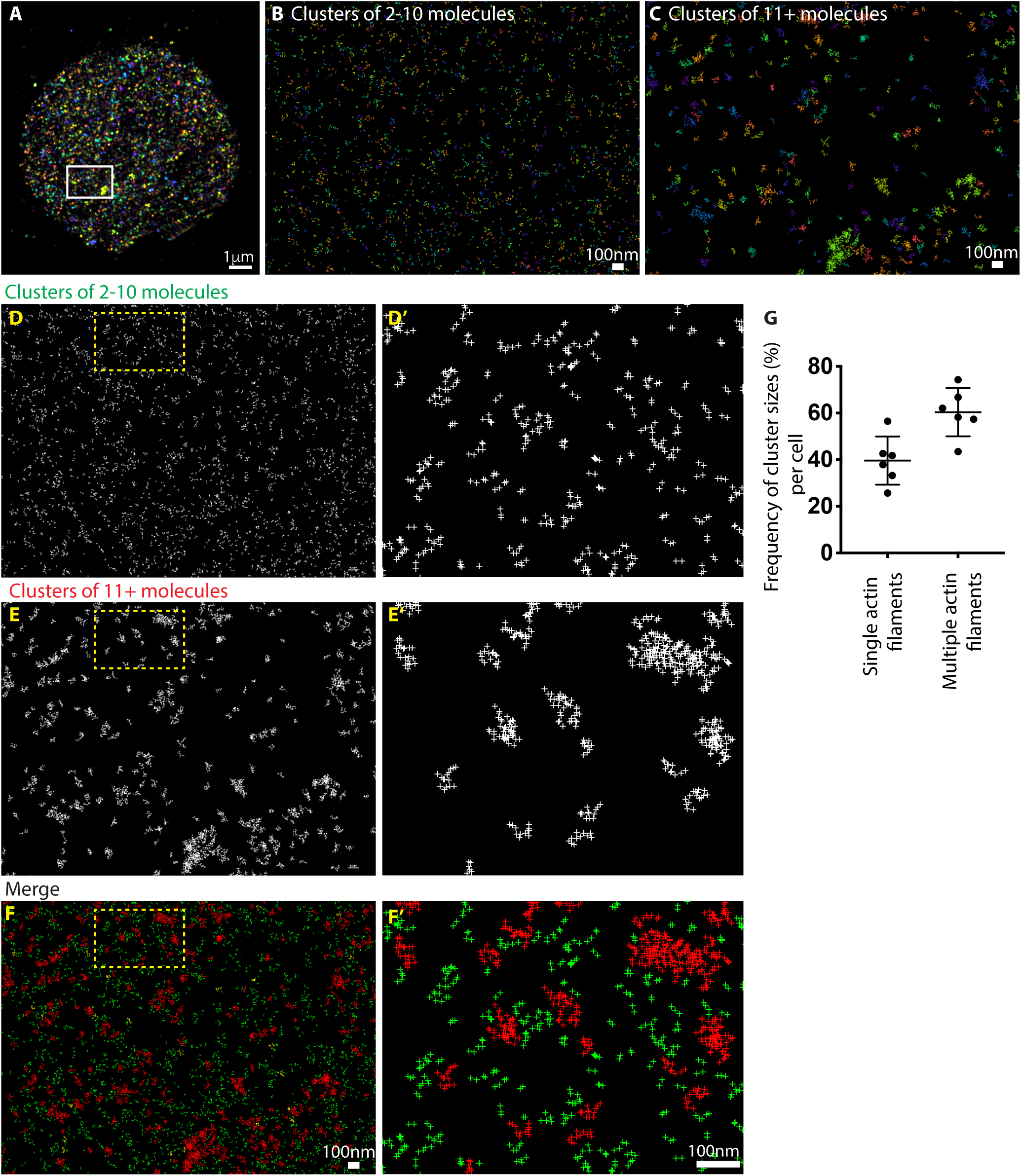
Visualization of locations of Alexa 647-phalloidin clusters containing 2-10 molecules or ≥11 molecules, from DBSCAN analysis of STORM images of RBCs. (A) DBSCAN analysis of STORM image of a RBC, with individual Alexa 647-phalloidin molecules color-coded to distinguish clusters (same cell as Figure 4B). White boxed region shown at higher magnification in panels B-F. (B) Visualization of clusters with 2-10 Alexa 647-phalloidin molecules, and (C) Visualization of clusters with ≥ 11 Alexa 647-phalloidin molecules, color coded as in B. Bars, 100 nm. (D, D’) Visualization of clusters of 2-10 molecules likely representing single short F-actins; yellow boxed region in D enlarged in D’. (E, E’) Visualization of clusters with ≥ 11 molecules likely representing groups of several or multiple short F-actins; yellow boxed region in E enlarged in E’. (F, F’) Merged images of clusters with 2-10 molecules (green) and clusters with ≥ 11 molecules (red); yellow boxed region in F enlarged in F’. Bars, 100 nm. (G) Frequency of clusters containing 2-10 molecules (Single actin filaments) or containing ≥ 11 molecules (Multiple actin filaments), per cell. Each point represents an individual cell. N = 6.

### Analyses reveal that non-uniform distribution of F-actins is not random

To test whether the irregular distribution of F-actin detected by STORM imaging and DBSCAN analysis could be explained by a random distribution of Alexa 647-phalloidins, we compared the experimentally determined distribution of Alexa 647-phalloidin molecules in STORM images with a randomly generated synthetic data set (Figure 6A, B). To generate the random data, we first calculated the number of Alexa 647-phalloidin molecules per cell and then distributed the same number of random points uniformly within the half of the surface area (∼65 µm^2^) of an RBC (Figure 6B). Then, we used DBSCAN with a minimum cluster size of 2 and a radius of 15 nm to compare clusters of different sizes in the randomly distributed molecules with the experimental data (Figure 6C, left panel). This analysis showed that in the case of randomly distributed molecules, almost 92% of the molecules are present in small clusters of 2-10 molecules and there are no clusters with a size larger than 20 molecules (Figure 6C, right panel). Thus, the frequency of single filaments (defined as clusters of 2-10 molecules) in the randomly generated synthetic data (∼90%) is significantly larger than that observed in the experimental data (∼40%), indicating that the experimentally observed distribution is not random.

**Figure 6.**
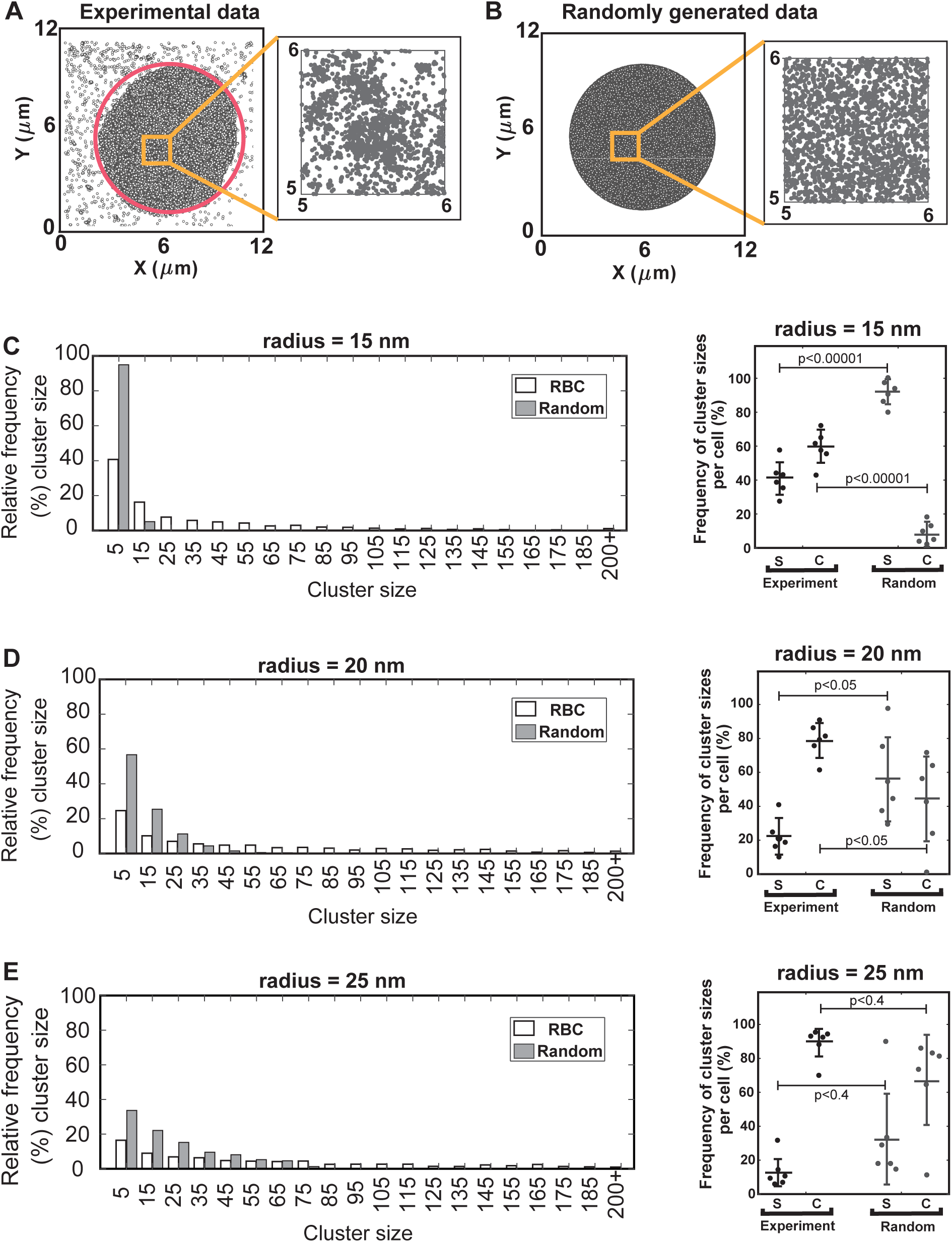
DBSCAN analysis of RBC STORM images versus randomly generated synthetic data reveals that a large proportion of the F-actins at the RBC membrane are distributed in groups of multiple filaments. (A) Scatter plot of distribution of Alexa 647-phalloidin molecules at the RBC membrane obtained from TIRF/STORM images as in Figure 4A. The red circle marks the RBC perimeter. Inset shows clustered distributions of Alexa 647-phalloidin molecules in the zoomed area. (B) Scatter plot of randomly generated uniform data distributed within the RBC area (red circle in panel A). The number of data points in random distributions is set equal to the number of Alexa 647-phalloidin molecules obtained from STORM images of RBCs in panel A. (C-E) Left panels, Histograms of Alexa 647-phalloidin cluster sizes and frequency in individual representative RBCs (open bars) compared to random data (shaded bars) for the same number of molecules for three different clustering radii of (C) 15 nm, (D) 20 nm and (E) 25 nm. A range of cluster sizes from 2-10 molecules up to > 200 molecules is observed, with larger clusters more abundant in experimental as compared to random data sets. Cluster sizes are grouped into bins of 2-10, 11-20, 21-30 etc., and centered on 5, 15, etc., for purposes of display. Right panels, Frequency of clusters with 2-10 Alexa 647-phalloidin molecules (single actin filaments, S) and clusters with ≥ 11 Alexa 647-phalloidin molecules (multiple actin filaments, C) in RBCs, versus random data for the three different clustering radii. The frequency of clusters with ≥ 11 Alexa 647-phalloidin molecules in RBCs, indicative of multiple actin filaments, is significantly larger than for the random data, with the exception of r = 25 nm, due to high variations among the random data sets at this radius. We report the frequency of cluster sizes for each distribution as Mean ± SD for N = 6 RBCs. Each point represents an individual RBC.

We also investigated the effects of varying the cluster radius on cluster size and frequency for both the experimental and synthetic data. Histograms of cluster size distributions showed that in the experimental data, the proportion of clusters with ≤10 molecules decreased while the proportion of clusters with ≥11 molecules increased as the clustering radius increased from 15 nm to 25 nm (Figure 6C-E). The same pattern was also evident in the random data with increasing radius, but the proportions of small clusters with ≤10 molecules in random data at all cluster radii was greater than for experimental data and the proportion of clusters with ≥11 molecules was lower, although this did not achieve statistical significance for r = 25 nm (Figure 6C-E). At a cluster radius of 15 nm, the frequency of clusters with ≥11 molecules in RBCs is almost seven times larger than a randomly generated uniform distribution (p<0.00001) (Figure 6C), while at a cluster radius of 20 nm, the frequency of clusters ≥11 molecules in RBCs is about two times greater than that of the random data (p<0.05) (Figure 6D). At a cluster radius of 25 nm, most molecules are in clusters for both the experimental and random data, so that the frequency of clusters with ≤10 molecules (single actin filaments) is ∼10% for experimental data and ∼30% for random data, but p<0.4 (Figure 6E). However, large variations in cluster frequency between individual random synthetic data sets for r = 25 nm make it difficult to compare the random data with the experimental data. We also evaluated the effect of setting the minimum number of molecules in a cluster to 3, for experimental and random data (Supplemental Figure 2). This analysis shows that the experimental data for a minimum cluster size of 3 demonstrates a greater frequency of larger cluster sizes compared to the random data (Supplemental Figure 2), similar to the data with a minimum cluster size of 2 (Figure 6). Therefore, we conclude that the F-actins are not uniformly distributed in a random fashion in RBCs, but rather that some filaments are present as solo filaments, which others form groups of multiple filaments, with a wide range of sizes on the RBC membrane.

## Discussion

### F-actin is distributed non-uniformly in the RBC membrane skeleton

Classic electron microscopy studies of the expanded RBC membrane skeleton have established its molecular organization as a regular 2D hexagonal lattice with F-actin nodes at vertices connected by long (α1β1)_2_–spectrin strands (Byers and Branton, 1985; Shen *et al*., 1986; Liu *et al*., 1987). Our observations here of nanoscale lattice organization in intact human RBCs indicate that the regular hexagonal pattern of F-actin nodes is not present in the unexpanded lattice *in situ*. Using TIRF and Zeiss Airyscan confocal microscopy of fluorescent-phalloidin-stained RBCs we discovered that F-actin was distributed non-uniformly across the entire cell, contrary to previous expectations (Pan *et al*., 2018), with numerous brightly stained F-actin foci upon a background of dimmer F-actin staining. A non-uniform actin filament distribution is not unique to human RBCs, as we had observed brightly stained F-actin foci in mouse RBCs imaged by TIRF microscopy (Sui *et al*., 2017). Further, STORM single molecule localization and DBSCAN analysis of human RBCs identified multiple filaments collected in groups of varying sizes, interspersed with single filaments, with the larger groups likely corresponding to the brightly stained F-actin foci and the single filaments and smaller groups to the dimmer background, respectively, detected by TIRF and Airyscan microscopy. Computational comparisons of STORM single molecule experimental data with synthetic data sets demonstrated that the experimentally observed non-uniform distribution of F-actins cannot be explained by a random uniform distribution of F-actins across the membrane.

Our data also do not support a uniform hexagonal arrangement of F-actin nodes in the lattice, ∼80 nm apart, where the (α1β1)_2_–spectrin strands are uniformly condensed ∼3-fold from their extended 200 nm length to accommodate the native membrane surface area, as depicted in Figure 7. This difference is readily apparent by visual comparison of a synthetic hexagonal distribution of Alexa 647-phalloidin clusters (Figure 7A) with an example of a TIRF/STORM single molecule image from our experimental data, showing an irregular distribution of F-actins (Figure 7B). Further, since 80 nm is below the resolution of TIRF and Zeiss Airyscan microscopy, a regular hexagonal distribution would have resulted in uniform F-actin staining by TIRF and Zeiss Airyscan, which we do not observe.

**Figure 7.**
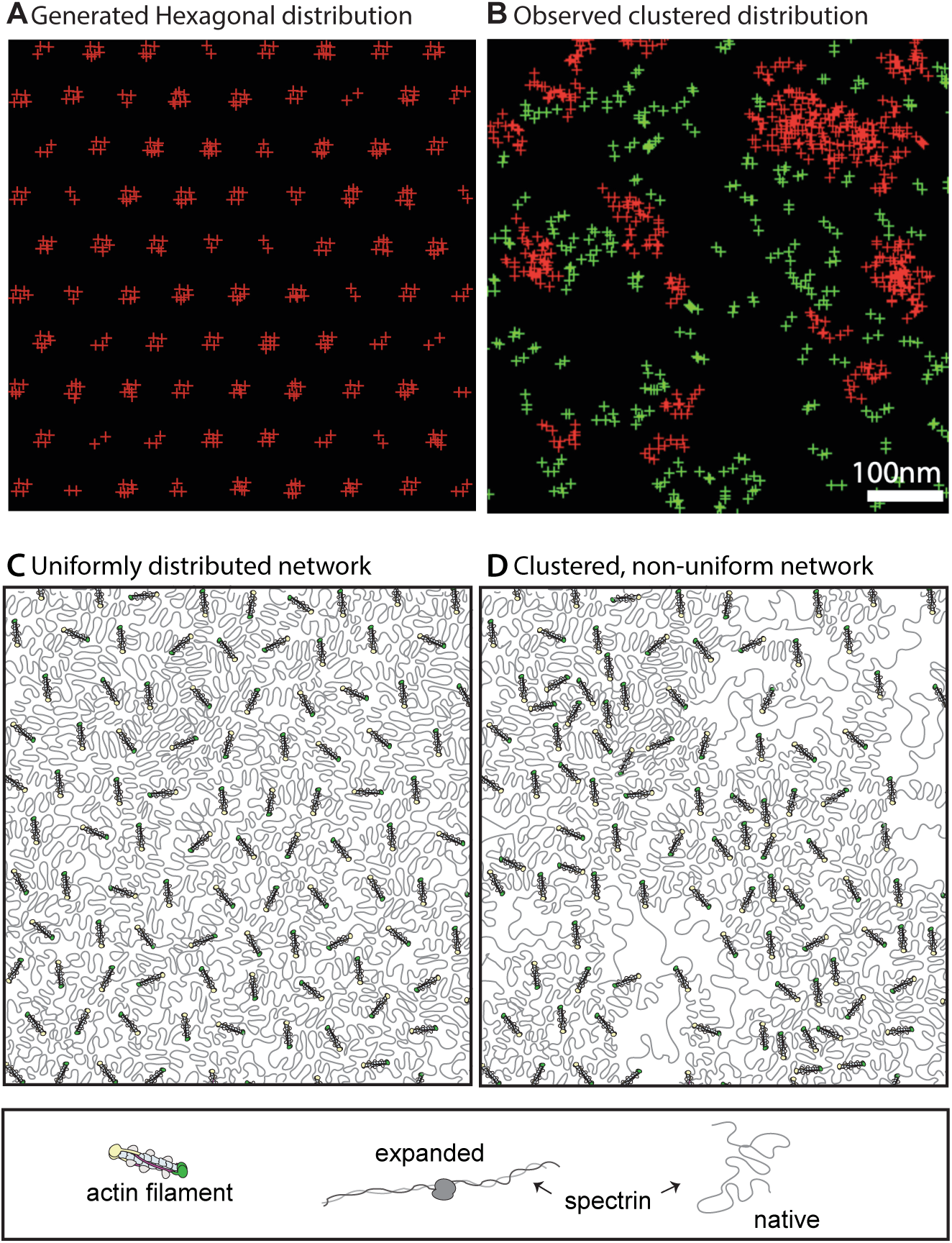
Comparison of a model with (A) hexagonally distributed clusters of Alexa 647-molecules ∼80 nm apart with (B) experimentally observed distribution of Alexa 647-molecules in the same sized region of an RBC. The hexagonal distribution in A was created manually in Keynote and B is from the image in Figure 5F’. Images are shown at the same scale. Bar, 100 nm. (C) Schematic of a spectrin-F-actin network with hexagonally distributed short F-actins connected by uniformly condensed spectrin strands, ∼80 nm long. (D) Schematic of a spectrin-F-actin network with irregularly distributed F-actins, with filaments in groups connected by shorter, condensed spectrin strands, and single filaments connected by longer, extended spectrin strands. Images and schematics are at the same scale.

The non-uniform distribution of F-actin nodes implies that the (α1β1)_2_–spectrin strands connecting the nodes are not all the same lengths, so that regions with F-actin foci (i.e., groups of F-actins) would presumably have connecting (α1β1)_2_–spectrin strands with shorter end-to-end lengths, while regions with sparser F-actins would have connecting (α1β1)_2_–spectrin strands with longer end-to-end lengths (Figure 7C-D). In support, electron microscopy of native and partially spread RBC membrane skeletons show that (α1β1)_2_–spectrin tetramers form a quaternary structure with variable degrees of extension, possibly due to mechanical tension (McGough and Josephs, 1990; Ursitti *et al*., 1991; Ursitti and Wade, 1993). Spectrin oligomer formation and lateral associations of spectrin strands observed in native membrane skeletons may also contribute to variations in spectrin end-to-end lengths between F-actins (Ursitti and Wade, 1993; Nans *et al*., 2011). Transient extension and condensation of spectrin strands, driven by thermal motion and or variations in mechanical tension on the membrane, could lead to transient F-actin rearrangements, explaining the dynamic fluctuations of F-actin foci observed by time-lapse imaging in live RBCs (Figure 3). However, it is not clear whether spectrin strand condensation and extension could be a driving force creating F-actin foci, or whether spectrin strands might change length in response to other forces driving F-actin associations (see below).

Since our observation of non-uniform F-actin organization was somewhat unexpected, we also performed a number of controls to rule out artefactual F-actin rearrangements due to fixation conditions, probes or imaging modality. Bright fluorescent-phalloidin stained F-actin foci were observed in intact flattened or biconcave RBCs fixed with a variety of methods, including paraformaldehyde, paraformaldehyde plus glutaraldehyde, or glutaraldehyde alone, and in cells labeled with either rhodamine phalloidin, Alexa 488-phalloidin, or Alexa 647-phalloidin. Moreover, F-actin foci were also observed in live, unfixed intact RBCs stained with the cell-permeable fluorescent analog of jasplakinolide, SiR-actin. A non-uniform distribution of F-actin was readily apparent in all microscopy modalities employed, including TIRF, Zeiss Airyscan, Leica SP8 Hyvolution, and TIRF/STORM.

What might explain the discrepancy of our observations with those of a recent super-resolution imaging study using STORM that revealed a uniform hexagonal lattice with F-actin nodes ∼80 nm apart (Figure 7A) (Pan *et al*., 2018)? One possibility is that the Fourier analyses demonstrating a hexagonal lattice organization were performed on relatively small regions of the membrane skeleton in this study, whereas we performed computational analyses of clustering on molecular localizations across the entire cell, determining the overall distribution of F-actin. Thus, we cannot exclude the presence of small regions of hexagonally organized F-actin nodes which would have gone undetected in our analysis. It is also possible that the bright F-actin foci/groups of filaments we observe here depend upon a population of filaments not detected in the previous study. For example, the fixation conditions used to prepare the cells for STORM imaging and computational analysis in the previous study may have led to preferential dispersion and/or disassembly of the clustered filaments.

It is tempting to use the known numbers of short F-actins per RBC (30-40,000) (Fowler, 2013; Lux, 2016) and the proportions of filament groups detected by STORM with DBSCAN to estimate the numbers of filaments in the bright F-actin foci detected by TIRF and Airyscan microscopy. TIRF detects ∼700 brightly stained F-actin foci per RBC, while Zeiss Airyscan confocal microscopy detects ∼1,680 foci per RBC, likely due to the greater sensitivity and higher resolution of Airyscan over TIRF microscopy. DBSCAN analysis of STORM single molecule Alexa-647 phalloidin localization data using a radius of 15 nm and cluster size of 2 indicates that ∼60% of molecules are in large clusters of varying sizes, with the remaining 40% as small clusters of <10 molecules, likely corresponding to actin subunits in single filaments (Figures 5-6). Further quantification indicates that 18 – 24,000 of the F-actins could be present in the foci detected by Airyscan imaging, so that on average, each of the ∼1,680 foci could contain between 11 – 14 individual F-actins (18 – 24,000 filaments/∼1,680 foci). However, this is likely to be a gross oversimplification since there is a ∼10-fold range in the STORM cluster sizes consisting of multiple filaments (Figures 5-6), and the relative fraction of multiple actin filaments *vs.* single filaments depends on both the clustering radius and minimum cluster size used for DBSCAN (Figure 6, Supplemental Figure S2). To evaluate the basis for the F-actin distribution, we focused on using random uniform distributions for the generation of synthetic data as the simplest way to test the nature of the F-actin distribution in experimental data. For a range of cluster radii and sizes, comparison of the synthetic data versus experimental data indicate that the distribution of F-actin in RBC is non-uniform. In addition, the precise number of actin subunits per 37 nm filament labeled with Alexa 647-phalloidin is uncertain. While labeling of RBCs with increasing concentrations of Alexa 647-phalloidin does not lead to brighter staining intensity or detection of additional molecular localizations in STORM (data not shown), it is possible that not all of the subunits in each filament may be available for phalloidin binding. However, despite this caveat and the computational uncertainties in the exact proportions of multiple *vs.* single filaments, all our data together clearly indicate an irregular distribution of F-actin in the RBC membrane skeleton.

### Molecular mechanisms to account for non-uniform distribution of F-actins

Dematin is an F-actin binding protein tightly associated with the short F-actins in the RBC membrane skeleton, present at a ratio of 1 dematin trimer for each short F-actin (Fowler, 2013). Dematin is a potent F-actin bundling protein that requires a so-called ‘headpiece’ domain to bundle F-actin, similar to villin, a bundling protein in intestinal epithelial cell microvilli (Siegel and Branton, 1985; Vardar *et al*., 2002). Another RBC F-actin binding protein, α,β-adducin, can also bundle F-actin *in vitro* (in addition to capping barbed filament ends) (Mische *et al*., 1987; Kuhlman *et al*., 1996). While mouse knockouts of dematin headpiece domain or *α*,*β−*adducin separately have a mild hemolytic anemia, their combined deletion results in a more severe anemia, with impaired membrane stability and network organization (Chen *et al*., 2007). Moreover, deletion of full-length dematin in mouse RBCs leads to a severe anemia due to loss of membrane stability with gross defects in spectrin-F-actin lattice organization (Lu *et al*., 2016). Based on our work here showing F-actin foci containing multiple filaments at the RBC membrane, it is attractive to speculate that cooperative F-actin bundling by dematin and/or α,β-adducin might explain formation of F-actin foci, and that reduced bundling in the absence of dematin or α,β-adducin might compromise lattice stability.

More generally, it is possible that the F-actins within the foci may represent a biochemically distinct population of F-actins, with a different complement of associated F-actin binding proteins. Interestingly, previous immunolocalization electron microscopy studies of dematin, tropomodulin1 (the RBC pointed end capping protein), *α*,*β* adducin (the RBC barbed end capping protein), and tropomyosin indicated that some but not all of the F-actin nodes were occupied by each of these F-actin binding proteins (Derick *et al*., 1992; Ursitti and Fowler, 1994). It is possible that such differences in occupancy could lead to differences in relative rates of actin assembly or disassembly that could contribute to dynamic rearrangements of F-actin foci and nanoscale irregular distributions of filaments (Gokhin *et al*., 2015; Gokhin and Fowler, 2016). Future studies will be required to co-localize dematin, α,β-adducin, tropomodulin1 and tropomyosin proteins with respect to F-actin foci in RBCs. Analysis of F-actin distribution and organization in RBCs from mice with genetic deletions of these F-actin binding proteins, using approaches described here, will also be informative.

Another potential mechanism that could explain formation of F-actin foci in the spectrin-F-actin lattice is cross-linking by nonmuscle myosin IIA (NMIIA). NMIIA forms bipolar filaments in RBCs that associate with F-actin in the membrane skeleton via their motor domains (Fowler *et al*., 1985; Smith *et al*., 2018). NMIIA motor activity promotes membrane tension and maintains RBC biconcave shape, controlling RBC deformability. Each NMIIA bipolar filament is ∼300 nm long, with ∼30 motor domains at each end (Smith *et al*., 2018; Pal *et al*., 2020), so that a NMIIA bipolar filament could bind to multiple F-actins simultaneously, creating foci. However, the ∼200 nm nearest neighbor distances of F-actin foci detected by Airyscan imaging (Figure 2D), and the diversity of Alexa-647 phalloidin cluster sizes detected by STORM (Figures 4-6), suggest that there is unlikely to be a direct correspondence between numbers of F-actins in foci and the F-actin binding motor domains at the ends of NMIIA bipolar filaments. Moreover, only a fraction of the F-actin foci would be able to interact with the NMIIA filaments at any one time, as there are ∼200 NMIIA filaments per RBC (Fowler *et al*., 1985), but ∼1,680 F-actin foci per RBC, based on ∼12 F-actin foci/μm^2^ detected by Zeiss Airyscan confocal microscopy (Figure 2D) and a 140 μm^2^ RBC surface area. If NMIIA does play a role in F-actin distributions, these filament associations would be expected to be transient, consistent with our observations of foci dynamics in time-lapse imaging of SiR-actin in live RBCs (Figure 3). This would also be consistent with our previous work showing that NMIIA associates with RBC F-actin via ATP-dependent interactions of the myosin motor domains (Smith *et al*., 2018).

Our previous computational modeling of force distributions in biconcave-shaped RBCs, together with immunostaining for NMIIA has shown that the forces exerted on the membrane and locations of NMIIA filaments are more concentrated at the dimple region as compared to the rim region (Alimohamadi *et al*., 2020). This implies that F-actin foci may be more abundant (denser), or more prominent (brighter), in the dimple as compared to the rim. However, with the approaches used here, we are unable to quantify F-actin foci in the dimple *vs.* the rim region of biconcave RBCs. In TIRF imaging, the dimple of a biconcave RBC is ∼1 μm from the coverslip and is not within the TIRF field. On the other hand, in the Zeiss Airyscan confocal images, due to the close proximity of dimple membranes from each side of the cell (∼0.5 μm), and the reduced resolution in Z (∼0.3-0.4 μm), *en face* images of F-actin foci at the membrane are difficult to ascribe to only one of the membrane surfaces (Figure 2C). Additional studies using other 3D high-resolution imaging approaches with fluorescent probes for both NMIIA and F-actin as well as investigation of F-actin clustering in RBCs treated with chemical inhibitors of NMIIA activity (Smith *et al*., 2018), or in RBCs with NMIIA mutations (Smith *et al*., 2019) will be necessary to further explore these ideas.

### Implications of nanoscale F-actin foci in the RBC membrane skeleton

The RBC membrane has long served as a model to elucidate membrane curvatures, protein interactions and dynamics that determine cellular signaling, shapes and movements in metazoan cells [for recent reviews, see (Kusumi *et al*., 2012a; Kusumi *et al*., 2012b; Fowler, 2013; Garcia-Parajo *et al*., 2014; Machnicka *et al*., 2014; Nicolson, 2014; Bennett and Lorenzo, 2016)]. Quantitative molecular models of forces to account for RBC biconcave shape and membrane deformations under flow or mechanical perturbations require precise knowledge of spectrin-F-actin lattice organization and interactions with the membrane bilayer. With the exception of a recent study (Feng *et al*., 2020), previous models have assumed a uniform distribution of F-actin nodes across the resting RBC membrane (Discher *et al*., 1998; Li *et al*., 2005; Li *et al*., 2007; Fedosov *et al*., 2010; Peng *et al*., 2013; Li and Lykotrafitis, 2014; Chen and Boyle, 2017; Fai *et al*., 2017). Notably, the recent study by Feng and colleagues (Feng *et al*., 2020) incorporates observed variations in spectrin strand length (Nans *et al*., 2011) along with increased measurements of spectrin density (Bryk and Wisniewski, 2017; Gautier *et al*., 2018) to provide a constitutive model of the membrane that more accurately reproduces the observed strain hardening behavior of the RBC membrane in experimental perturbations (Feng *et al*., 2020).

Studies of membrane protein mobility were pioneered in RBCs where mobility was shown to be restricted by direct anchorage of transmembrane proteins to the spectrin-F-actin network, or by trapping within ‘corrals’ created by the extended spectrin tetramers of the lattice (Fowler and Branton, 1977; Fowler and Bennett, 1978; Golan and Veatch, 1980; Sheetz, 1983; Kodippili *et al*., 2012). Quantitative analyses of membrane protein mobility in RBCs have generally presumed a uniform network, with ‘corral’ dimensions corresponding to a triangular lattice of extended spectrin tetramers linked to uniformly distributed F-actin nodes (Kodippili *et al*., 2020). Our data here indicate that mechanisms of RBC membrane protein mobility will require reevaluation, accounting for considerable heterogeneity in corral dimensions, based on our data showing closely spaced nanoscale F-actin clusters interspersed with individual filaments further apart. Indeed, nanoscale clustering of membrane proteins has emerged as a prevalent feature of membrane protein dynamics and plasma membrane organization in other cells (Kusumi *et al*., 2012a; Kusumi *et al*., 2012b; Garcia-Parajo *et al*., 2014; Nicolson, 2014; Bennett and Lorenzo, 2016).

The irregular distribution of F-actin in native RBCs appears to contrast with neurons, where super-resolution fluorescence microscopy reveals a regular 1D periodic lattice with circumferential rings of F-actin connected by extended (*α*_2_*β*_2_)_2_-spectrin tetramers ∼190 nm long, along the length of axons and dendrites (Xu *et al*., 2013; Zhong *et al*., 2014; D’Este *et al*., 2015; Leterrier *et al*., 2015; Leterrier, 2021). However, it is attractive to consider that the distribution of the short F-actins within the rings themselves may correspond to the nanoscale F-actin foci we observe in RBCs. This idea is supported by the presence of dematin, tropomyosin3.1 and the actin capping proteins, tropomodulin1 and *αβ*-adducin, within the axonal F-actin rings (Abouelezz *et al*., 2020; Zhou *et al*., 2020), indicating that the rings could consist of many short filaments distributed in an elongated “cluster” spanning the axon circumference. On the other hand, a recent study using platinum replica EM to visualize F-actin in mechanically unroofed axons suggests that rings may consist of long, braided F-actins (Vassilopoulos *et al*., 2019). Since the same sets of actin binding proteins are present in the RBC spectrin-F-actin network, and in axonal F-actin rings, could RBC F-actin foci contain longer filaments as well? Future ultrastructural analyses of the RBC spectrin-F-actin network will be required to answer this question.

Nanoscale F-actin foci are also likely to be characteristic of spectrin-F-actin networks in non-erythroid cells. A recent study from the Gauthier lab shows that large micron-sized spectrin-rich domains on the plasma membrane are located between F-actin-rich stress fibers in mouse embryonic fibroblasts (Ghisleni *et al*., 2020). While the spectrin-rich domains are F-actin-poor with respect to the stress fibers, closer inspection reveals F-actin staining in the spectrin-rich regions visualized by TIRF microscopy. Strikingly, both the F-actin and the spectrin staining intensity appear non-uniform in these regions, resembling the non-uniform F-actin distribution we observe at the RBC plasma membrane. Inasmuch as spectrin-rich domains restrict clathrin-mediated endocytosis (Jenkins *et al*., 2015; Ghisleni *et al*., 2020), further investigation of the nanoscale distribution and dynamics of spectrin and F-actin in these plasma membrane regions and how it may preclude or convert to one favoring endocytosis will be a fascinating area of future investigation.

In conclusion, we expect that the relative simplicity of RBC model system will continue to provide an opportunity to scrutinize the molecular and structural basis for membrane protein organization and dynamics, and the role of spectrin-F-actin networks in determination of membrane functional properties.

## Materials and Methods

### Fluorescence staining of RBCs

Whole blood was collected from healthy human donors into EDTA tubes (BD Diagnostics) by the Normal Blood Donor Service at The Scripps Research Institute, La Jolla, CA. For phalloidin staining, 20 μl of whole blood was added to 1 ml of 4% paraformaldehyde (PFA, Electron Microscopy Sciences) in Dulbecco’s PBS (DPBS – Gibco), mixed, and incubated overnight at room temperature, unless otherwise indicated. In some experiments, cells were fixed in 4% PFA for 4h at room temperature, or in a mixture of 4% PFA and 0.01% glutaraldehyde (Electron Microscopy Sciences) in DPBS for 30 min at room temperature, followed by washing and incubation with 0.1% sodium borohydride in DPBS for 20 minutes at room temperature. Fixed RBCs were washed three times in DPBS by centrifuging for 5 minutes at 1000 × *g*, permeabilized in DPBS + 0.3% TX-100 for 10 minutes, and then blocked in 4% BSA, 1% normal goat serum in DPBS (Blocking Buffer, BB) and kept in blocking buffer at 4°C for up to one week. 20μl of fixed, permeabilized and blocked RBCs plus 180μl of blocking buffer were then incubated with fluorescent-phalloidin and fluorescent-conjugated glycophorin A (GPA) or fluorescent-conjugated wheat germ agglutinin (WGA) for 1-2 h at room temperature, followed by washing three times in BB as above. Fluorescent phalloidins were rhodamine-phalloidin (Life Technologies R415 at a final concentration of 130nM), or Alexa-488 phalloidin (Life Technologies). Fluorescent-GPA was Alexa 488-GPA (BD Biosciences GA-R2 1:100) and Alexa-555-WGA (Molecular Probes 2ug/ml). Stained cells were cytospun (Thermo Scientific Cytospin 4 1000rpm for 3 minutes) onto 1.5 thickness coverslips and mounted with ProLong^TM^ Gold mounting medium (Molecular Probes #P36934) onto slides prior to imaging. For live cell microscopy, cells in 20μl of whole blood from a normal donor were washed twice in 1.3mL of DPBS/0.05%BSA. Washed cells were diluted 1:9 and stained with 50 nM SiR-actin (Spirochrome/Cytoskeleton) for 30 minutes (Lukinavicius *et al*., 2013). Stained cells were settled on Mat-tek dishes (MatTek, Ashland, MA) with a 1.5 size coverslip bottom for imaging.

### Fluorescence Microscopy

Live or fixed fluorescently stained RBCs were imaged using a Nikon Ti inverted microscope with a 100× Apochromat oil objective (NA 1.49), either by epifluorescence microscopy using vertical illumination with 488 and 561 laser lines and an ORCA-Flash 4.0 V2 Digital CMOS camera (Hamamatsu) or by TIRF illumination with the same laser lines, and DIC microscopy. Images were acquired using NIS-Elements 5.0 software. (Figure 1). Fixed RBCs were also imaged using a sensitive Zeiss LSM 880 Airyscan laser scanning confocal microscope with a 63× or 100× 1.46 NA oil Plan Apo objective (Figure 3). Z-stacks were acquired at a digital zoom of 1.7 and a Z-step size of 0.168 μm (Figure 2). Note that standard confocal laser scanning microscopy was unable to image fluorescent-phalloidin staining in RBCs due to the dim signal, as a consequence of quenching from the abundant hemoglobin (Smith *et al*., 2018), relatively low sensitivity of standard confocal microscopy, and photobleaching during confocal image acquisition. Live cell images of SiR-actin labeled RBCs were also acquired using a Leica TCS SP8 STED 3X microscope, equipped with a HC PL APO CS2 93x/1.30 glycerin objective lens. A 3D time series was collected with a detector window set to collect fluorescence emission at 655 - 751 nm. A white light laser was used to excite the fluors. STED images were collected using depletion lasers of 592, 660, 775 nm, HyD detectors, and time gating of fluorescence emission between 0.3 – 6.5 ns. Bidirectional scanning at a frequency of 8,000 Hz, with a scan resolution of 512 x 512 pixels and a line averaging of 5. Z stacks were collected with a step size of 0.333 µm, every 2.8 seconds. Post-acquisition, the images were deconvolved using SVI Huygens Essential software.

### Image analysis for TIRF and Zeiss AiryScan images

TIRF images were deconvolved using the 2D deconvolution module in Nikon Elements, NIS-Elements 5.0 software. The General Analysis tool in Elements was used to pinpoint centroids in the segmented deconvolved images, and to calculate the nearest neighbor distances between the spots. AiryScan confocal stacks of overnight-fixed RBCs stained with GPA or WGA for membrane, and phalloidin for F-actin were processed using Zen 2.3 software and analyzed using Volocity 6.3.0 software. The outer 4-6 z-slices of the Airyscan stacks, corresponding to the outer 300-500nm of each cell, were cropped in Nikon Elements. Extended focus projections of these stacks were exported into Tiff images and spot identification and density (spot/μm^2^) measurements were made using Elements software the same way as for TIRF images. For Figure 2C, pixel intensity measurements of the WGA and phalloidin stains were performed using similar thresholding parameters in the find object option in the measurement module of Volocity 6.3.0. Coefficients of variation were calculated from the standard deviation and mean intensity values for each stain.

### STORM imaging of F-actin

STORM imaging was performed as in (Xu *et al*., 2013; Pan *et al*., 2018), with minor modifications. Briefly, whole blood was obtained as above and Dulbecco’s PBS (DPBS – Gibco [2.67 mM KCl, 1.47 mM KH_2_PO_4_, 137.93 mM NaCl, 8.06 mM Na_2_HPO_4_-7H_2_O]) containing 10 mM glucose and 5 mg/mL BSA. Cells were washed twice in this buffer by centrifugation at 1200 rpm for 5 min in a Beckman Avanti centrifuge swinging bucket rotor and resuspended at 3.0x10^6^ cells in PBS with 10 mM glucose and 5 mg/mL BSA. RBCs were allowed to settle onto poly-L-lysine coated coverslips (squeaky clean coverslips were coated with a 0.1% (w/v) solution of poly-L-lysine (Sigma-Aldrich P8920) 1-2 h room temp with poly-L-Lysine then dried and kept at 4°C until ready for use), for 10 min at room temperature (1.5 x 10^6^ cells per coverslip). Attached cells were permeabilized in 0.0015% saponin in cytoskeleton buffer (CB; 10 mM MES, pH 6.1, 150 mM NaCl, 5 mM EGTA, 5 mM glucose, and 5 mM MgCl_2_) (Xu *et al*., 2013; Pan *et al*., 2018) for 4 min at room temperature, fixed in either 2% glutaraldehyde or 4% paraformaldehyde (Electron Microscopy Sciences) in CB for 20 min room temperature, and reduced with freshly prepared 0.1% sodium borohydride in PBS for 20 min at room temperature. Cells were stained with 0.4 µM Alexa 647-Phalloidin (Life Technologies) for 40 min in 3% BSA in PBS, washed in 3% BSA in DPBS, mounted with ∼ 20μl Vectashield (Vector Laboratories Cat. No. H-1000) and sealed with nail polish. STORM imaging was performed on a Nikon Ti inverted microscope with a 100× Apochromat oil objective (NA 1.49), using TIRF illumination with the 647 nm laser line and an ORCA-Flash 4.0 V2 Digital CMOS camera (Hamamatsu) (Figure 4). 100,000 images, 10 ms each, were acquired using the STORM acquisition module in NIS-Elements 5.0 software. STORM images were processed for peak height of 175 with drift correction and analyzed using the DBSCAN function embedded in Nikon Elements (Figure 4). Data was exported to excel for statistical analysis and graph plotting in Prism 8.2 software.

### DBSCAN clustering analysis of experimental *vs.* synthetic datasets

We used the DBSCAN algorithm (Schubert *et al*., 2017) implemented in the python library scikit-learn (Pedregosa *et al*., 2011) for the clustering tasks with parameters epsilon (eps) and min_samples. eps denotes the maximum distance between two samples to be considered as the neighborhood of each other. In this study, we did the clustering analysis for three different values of eps, eps=15 nm, eps=20 nm, and eps=25 nm. The min_samples indicates the number of samples in a neighborhood for a point to be considered as a core point, including itself. Here, we ran the algorithm for both min_samples=2 (Figure 6) and min_samples=3 (Supplemental Figure 2). After clustering the data, the algorithm gathers statistics on the results of the clusters, including the number of molecules per cluster size and the number of outliers, which are the molecules that are too far from any other molecules.

### Statistical analysis

Data are presented in dot plots as mean ± standard deviation (SD). Differences between means were detected using unpaired t-tests. When more than one comparison was made, means were compared using one-way ANOVAs followed by Tukey’s multiple comparisons test. Statistical significance was defined as p < 0.05. Statistical analysis was performed using GraphPad Prism 7.03 software. Box and whisker plots show median values (horizontal center line), third and first quartiles (top and bottom of boxes), and the minimum and maximum of the data (whiskers), as well as the mean (+ sign).

## Acknowledgements

We gratefully acknowledge Scott Henderson and the Scripps Microscopy Core for assistance with Leica SP8 Hyvolution microscopy of live RBCs. This work was supported by National Institutes of Health/National Heart, Lung and Blood Institute Grant R01-HL083464 (to V.M.F.) and National Institutes of Health/National Institute of General Medicine Grant R01-GM132106 (to P.R.). H.A. gratefully acknowledges partial support from a Siebel Scholarship.

---

**Supplemental Figure S1.**
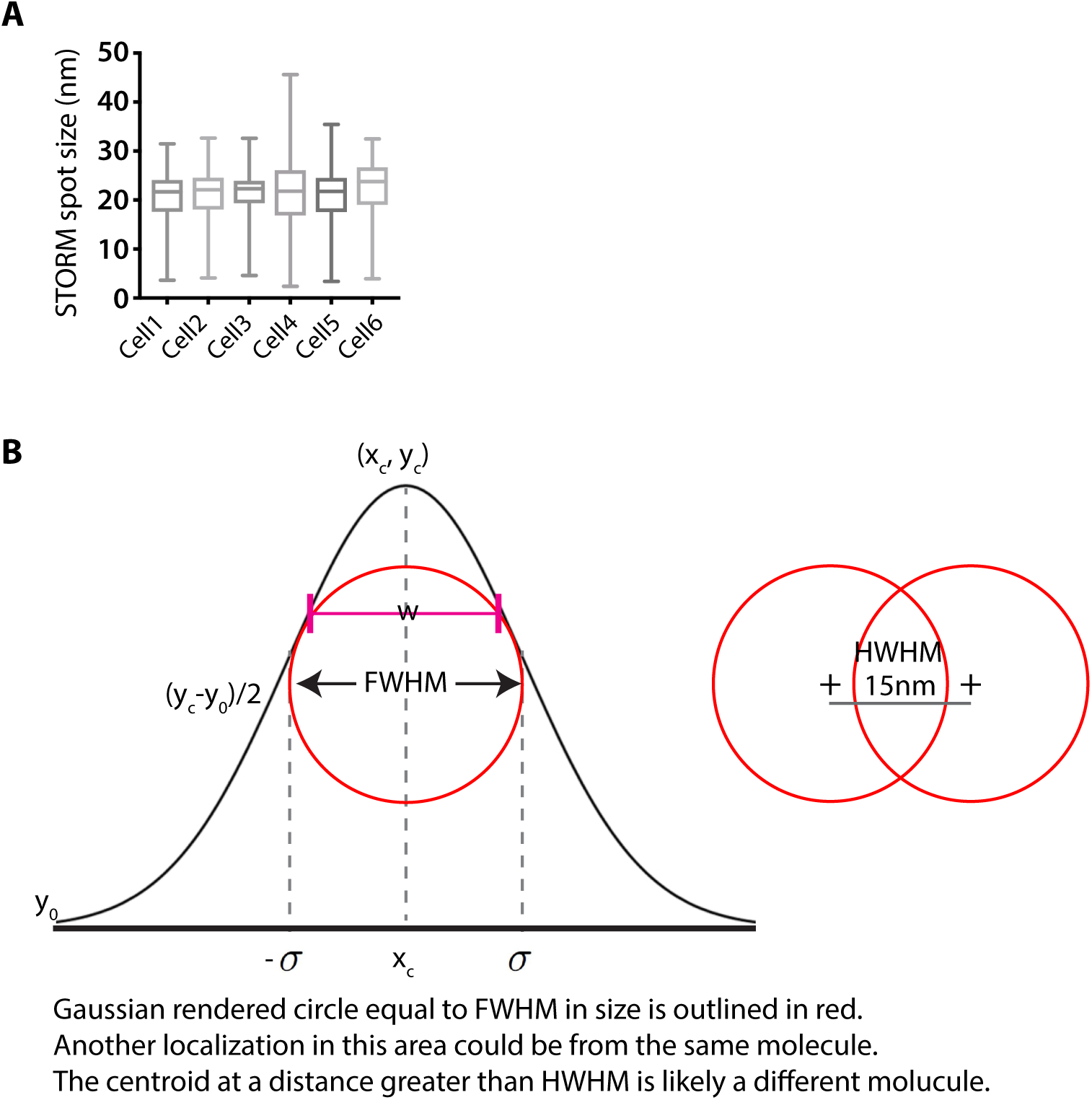
(A) STORM spot sizes for Alexa 647-phalloidin molecules in each of 6 individual RBCs. The average value for each cell was Cell 1, 20 +/- 4 nm, n = 101,139; Cell 2, 21 +/- 4 nm, n = 107,836; Cell 3, 21 +/- 4 nm, n = 45,800; Cell 4, 21 +/- 5 nm, n = 128,838; Cell 5, 21 +/- 5 nm, n = 121,983; Cell 6, 22 +/- 5 nm, n = 117,846. The line in the box indicates the median, and the box shows the upper 25% and lower 75% and the whiskers the minimum and maximum. (B) Left, Schematic of Gaussian intensity distribution for a fluorescence spot, showing the Full Width Half-Maximum (FWHM) of the signal at 2.355 σ (where σ is standard deviation). Right, The minimum detectable distance between two fluorescent spots is considered to be ½ FWHM, ie, Half Width Half-Maximum (HWHM). With a spot size of 20.68 nm +/- 3.77 nm to 22.63 nm +/- 5.83 nm (S.D.) for individual cells (Panel A), and FWHM = 2.355 σ, we calculated the HWHM to be between 8.88 nm and 13.73 nm. The centroids of fluorescence signals ≥ 15 nm apart are considered to be from two individual molecules, while signals < 15 nm apart are considered to be from the same molecule.

**Supplemental Figure S2.**
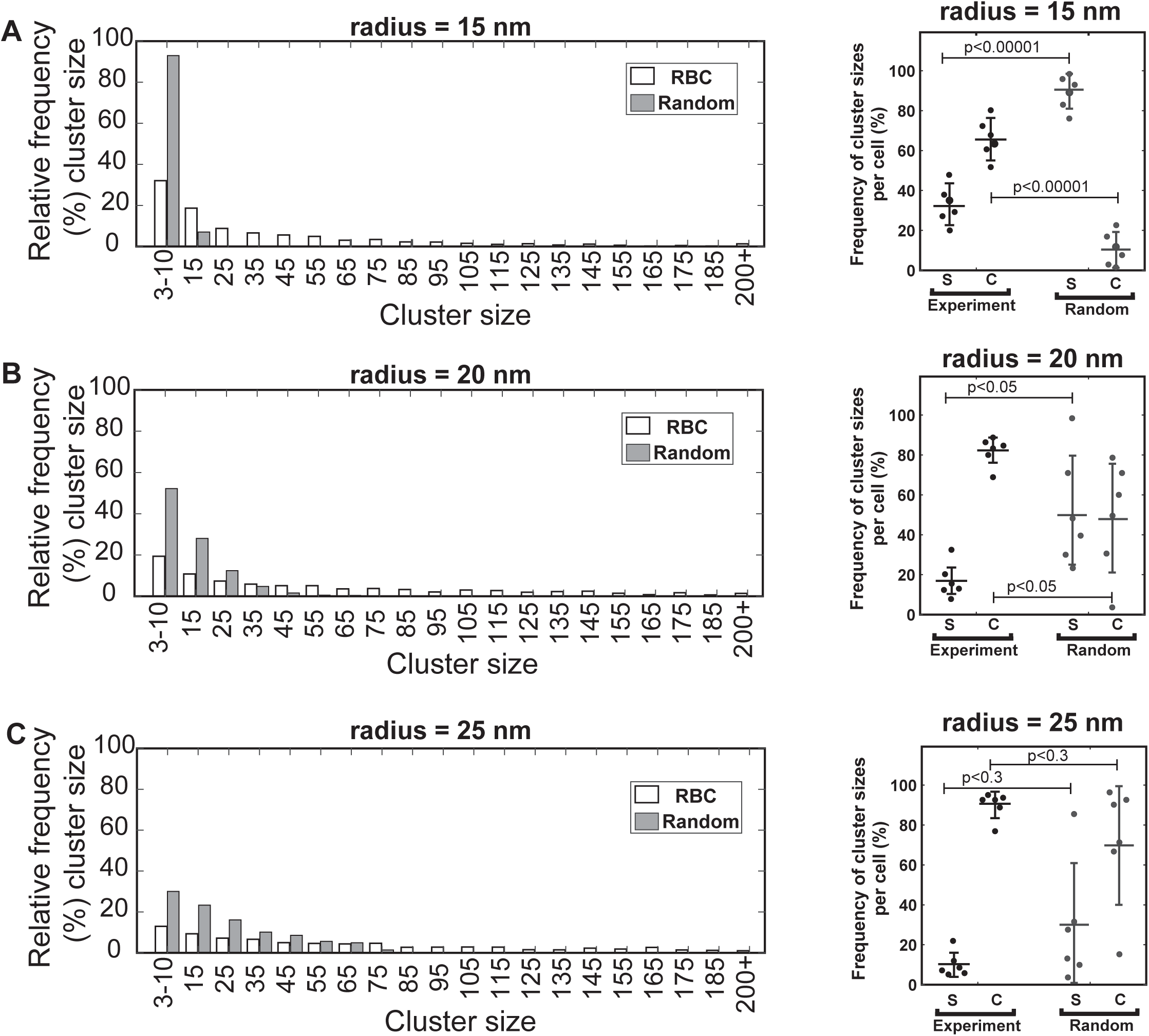
DBSCAN analysis using 3 points minimum per cluster for experimental RBC STORM images versus randomly generated data. (A-C) Left Panels, Histograms of Alexa 647-phalloidin cluster sizes and frequency in individual representative RBCs (open bars) compared to random data (shaded bars) for the same number of molecules for three different clustering radii of (C) 15 nm, (D) 20 nm and (E) 25 nm. A range of cluster sizes from 3-10 molecules up to > 200 molecules is observed, with larger clusters more abundant in experimental as compared to random data sets. At r = 15 nm for random data, almost 90% of molecules are in clusters of 3-10, and there are no clusters with a size > 20 compared to RBCs where only 30% are in clusters of 3-10 and most molecules are in clusters with size ≥ 11. For r = 20 or 25 nm, the random data also contains greater frequencies of smaller clusters as compared to RBC data. Cluster sizes are grouped into bins of 3-10, 11-20, 21-30 etc., and centered on 5, 15, etc., for purposes of display. Right panels, Frequency of clusters with 3-10 Alexa 647-phalloidin molecules (single actin filaments, S) and clusters with ≥ 11 Alexa 647-phalloidin molecules (multiple actin filaments, C) in RBCs versus random data for the three different clustering radii. The frequency of multiple actin filaments in RBCs is significantly larger than in the random data, with the exception of r = 25 nm, due to high variations among the different random data sets at this radius. We report the frequency of cluster sizes for each distribution as Mean ± SD for N = 6 RBCs. Each point represents an individual RBC.

